# Proteome profiling of plant Cajal bodies uncovers kingdom-specific components

**DOI:** 10.1101/2025.07.10.664108

**Authors:** Yu Zhou, Huang Tan, Angela Vicente-Luque, Marvin Weis, Lena Wehinger, Marina Schreidl, Irina Droste-Borel, Boris Macek, Kenneth Wayne Berendzen, Rosa Lozano-Durán

## Abstract

Insight into the molecular composition of cellular microenvironments is essential for understanding their function. Membraneless organelles (MLO), such as the nuclear Cajal body (CB), play fundamental roles in eukaryotic cells, but their very nature poses a challenge to biochemical isolation. Despite the relevance of the CB for cell function in plants and animals, its proteome remains to be profiled. Here, we purify plant CBs using a combination of biochemical isolation and fluorescence-activated particle sorting (FAPS) and define their proteome by mass spectrometry. Our work identifies 15 and 95 proteins uniquely localized and enriched in CBs, respectively, of which 22 are also labelled by CB markers in proximity labelling assays. CB proteins are preferentially involved in RNA processing, chromatin organization, and gene expression. While a partial overlap with known animal CB proteins exists, our data reveal a distinct, kingdom-specific CB proteome. These results provide the first comprehensive proteome of CBs and establishes a foundation for dissecting the molecular functions of this organelle across multicellular lineages.

## MAIN TEXT

Heterogeneity and compartmentalization are fundamental features of living cells, with cellular activity spatially distributed across different microenvironments. Beyond the hallmark eukaryotic membrane-bound organelles, membraneless organelles (MLOs) and biomolecular condensates have emerged in recent years as widespread cellular structures with essential biological roles. Nevertheless, the full functionality of such molecular assemblies remains in most cases to be determined. A first necessary step towards a complete understanding of these structures is the definition of their molecular components; this is however challenged by their dynamic nature, their potential compositional complexity and, importantly, the difficulties in their isolation.

One such MLO is the Cajal body (CB). The CB is a nuclear condensate with a diameter of 0.15-2 µm, initially described in vertebrate neurons by Santiago Ramón y Cajal in 1903, found in animals and plants and believed to form via liquid-liquid phase separation (LLPS). These bodies are present in varying numbers per nucleus, form at the sites of transcription and processing of small nuclear/nucleolar RNAs (SnRNA/SnoRNA), and play a role in the biogenesis and assembly of ribonuclear protein (RNP) particles^1^. The defining component of the CB is coilin, a conserved scaffolding protein required for formation of this MLO^2^; coilin undergoes LLPS *in vivo* through its intrinsically disordered region (IDR) domain^3^. Beyond coilin, however, our knowledge on CB composition is limited. Recent studies have characterized the proximal proteome of coilin by proximity labelling (PL) in animal cell cultures^4–6^. Nevertheless, the resulting dataset is likely only a partial CB proteome, restricted to proteins in a certain spatial domain. The characterization of the complete CB proteome would require purification of this organelle, a non-trivial task considering its membraneless nature.

Strikingly, the compositional information on the plant CB is even more scarce. Filling this knowledge gap is particularly pressing given the relevance of CBs for plant biology^7^ and the hints at potential fundamental differences between animal and plant CBs. For example, plant CBs harbour components of the plant-specific RNA-directed DNA methylation (RdDM) pathway^8^, and have emerged as convergent targets of viruses^7,9^. On top of that, some of the coilin proximal proteins identified in human cell cultures lack obvious orthologs in plants^4,5^.

To investigate the protein composition of plant CBs, we developed a method based on fluorescence-activated particle sorting (FAPS) to isolate this MLO from *Nicotiana benthamiana* leaves (Figure 1A). To enable specific labelling of CBs, we fused the CB scaffold protein NbcoilinA^10^ to GFP and expressed it under the control of the ribosomal protein subunit 5a (pRPS5A) promoter^11^. The resulting NbcoilinA-GFP fusion protein localized specifically to CBs, as indicated by co-localization with another CB marker, AtU2B″^12^ (Figure 1B).

**Figure 1.**
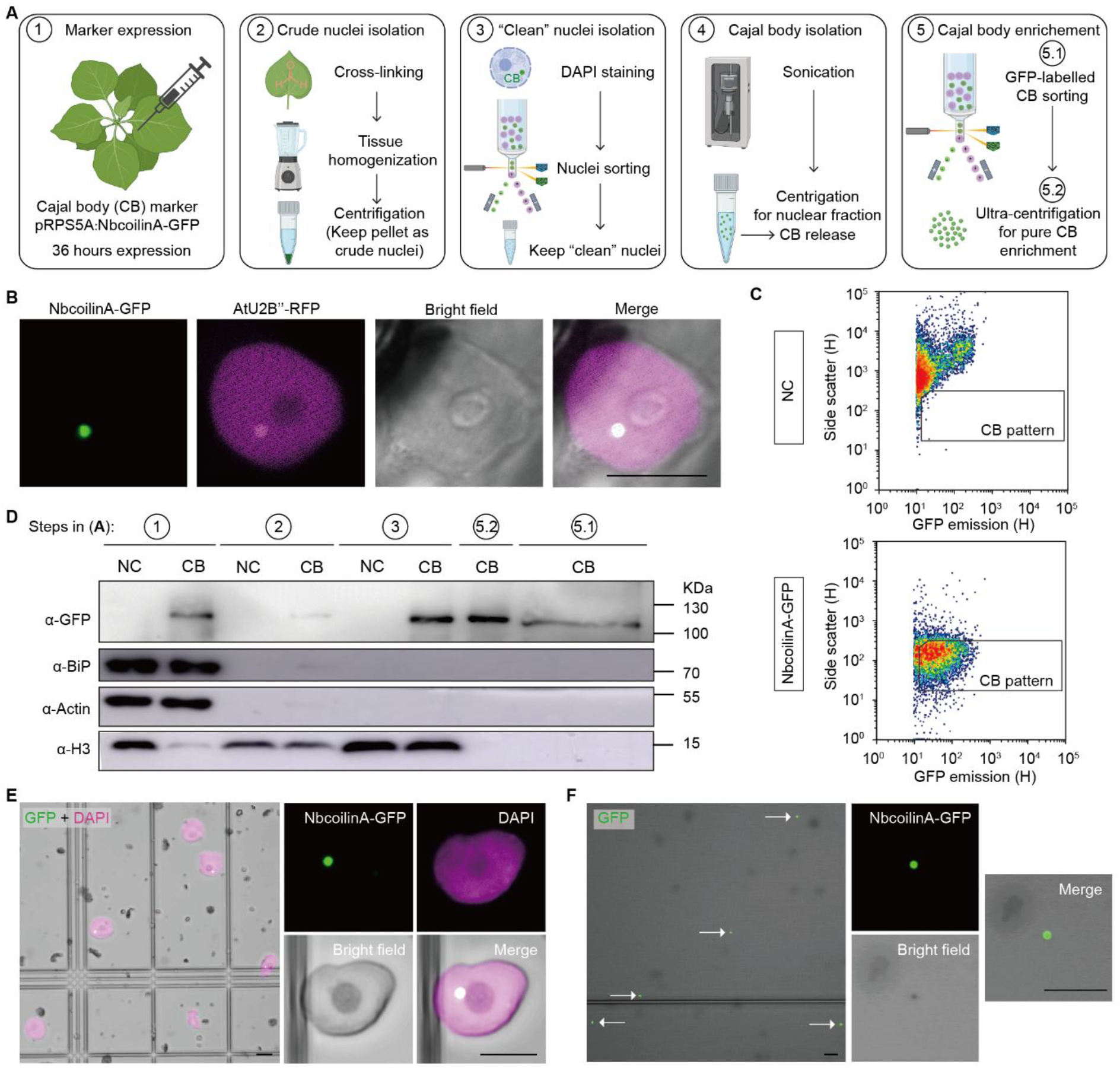
Isolation of Cajal bodies from *N. benthamiana* using fluorescence-activated particle sorting. A. Schematic overview of the Cajal body (CB) isolation workflow. Leaves expressing a CB-marker (NbcoilinA-GFP) were crosslinked with formaldehyde. Crude nuclei were extracted, stained with DAPI, and sorted. Following sonication and ultracentrifugation, GFP-positive particles were sorted to enrich for CBs. Leaves transformed with an empty vector were used as negative control (NC). B. Confocal imaging showing the specific CB localization of NbcoilinA-GFP; AtU2B’’-RFP is used as CB marker. Scale bar = 10 µm. C. Representative flow cytometry plots of side scatter versus GFP emission used to define the CB gating strategy. GFP-positive particles present in the marker-expressing samples but absent in the NC were selected. H: height. D. Immunoblot analysis of samples from each purification step from A, showing enrichment of NbcoilinA-GFP (∼120 kDa) and absence of major organelle contaminants. Markers include BiP (Endoplasmic reticulum, 80 kDa), Actin (cytosol, 45 kDa), and Histone H3 (nucleus, 17 kDa). E. Confocal images of sorted nuclei (“clean nuclei”) stained with DAPI (magenta) and expressing NbcoilinA-GFP (green). Scale bars = 10 µm. F. Confocal images of GFP-positive particles representing FAPS-isolated CBs. Arrowheads mark CB. Scale bars = 10 µm.

NbcoilinA-GFP was transiently expressed in fully expanded leaves of four-week-old *N. benthamiana*. As a negative control for the entire isolation procedure, an empty vector lacking the GFP tag (NC) was used in parallel. Transient transformation with the empty vector resulted in no detectable GFP signal and was used to define the background for the isolation. Samples were harvested at 36 hours post-infiltration (hpi) and cross-linked with formaldehyde to preserve CB integrity during isolation. The tissue was further homogenized using a kitchen blender to release crude nuclei, which were then enriched by fluorescence-activated nuclei sorting (FANS) following 4′,6-diamidino-2-phenylindole (DAPI) staining. FANS was initiated by setting a fluorescence threshold for DAPI and applying a side scatter (SSC) gate (Supplementary Figure 1A). Both nuclei from NbcoilinA-GFP and NC samples were identified as DAPI-positive populations with distinct scatter profiles. To confirm the presence of GFP-positive nuclei in the NbcoilinA-GFP sample, DAPI fluorescence was plotted against GFP fluorescence. In contrast to the NC samples, which lack GFP expression, the NbcoilinA-GFP sample displayed a distinct GFP-positive nuclei population.

The purified nuclei were disrupted by sonication to generate a nuclear extract, from which intact CBs were isolated using FAPS. GFP-positive particles were identified as small structures with lower side scatter intensity than nuclei during FAPS (Figure 1C), consistent with the expected size and complexity of CBs. Gating parameters were optimized by comparing GFP versus SSC plots across NbcoilinA-GFP and NC. This gating strategy enabled the reproducible isolation of a specific and consistent population of GFP-positive particles in NbcoilinA-GFP expressing samples, which was absent in the NC. To further confirm that the FAPS-isolated particles were enriched in NbcoilinA-GFP, an immunoblot was performed using an anti-GFP antibody on equal amounts of protein from leaf tissue, crude nuclei, purified nuclei, and FAPS-isolated particles from samples expressing the NbcoilinA-GFP or transformed with the NC (Figure 1D). A clear GFP signal was detected in the whole leaf, crude nuclei, purified nuclei, and FAPS-sorted particles from NbcoilinA-GFP expressing samples but was absent in all corresponding NC samples. Notably, the isolated GFP-positive particles could be recovered by ultracentrifugation, and the GFP signal intensity in these fractions was comparable to that of purified nuclei. In contrast, the nuclear protein histone H3 was significantly depleted in the isolated particles but enriched in the “clean” nuclei sample, indicating that the method specifically enriches for NbcoilinA-GFP-positive particles. To evaluate contamination from other organelles, immunoblots were also performed using anti-actin and anti-BiP antibodies, recognizing cytoplasm and endoplasmic reticulum (ER) proteins, respectively. Both markers were enriched in leaf samples but barely detectable in nuclei or FAPS-sorted particles. These results indicate that the isolation protocol effectively removes cytoplasmic and ER contaminants, yielding highly purified CB-like isolations. Confocal microscopy confirmed that FANS purified nuclei remained intact, and the GFP signal was unaltered, indicating that the structure of CBs was successfully preserved (Figure 1E) as following FAPS, small round GFP-positive particles, consistent with the expected size and morphology of CBs, were observed (Figure 1F).

CB-like GFP-positive particles isolated from *N. benthamiana* leaves were subjected to LC-MS/MS analysis, alongside nuclear fractions from the same samples (input) and from NC (see Methods for details). Label-free quantification (LFQ) values from the input and NC fractions were used as background to identify proteins specifically enriched in the isolated CBs following the filtering method described in Figure 2A.

**Figure 2.**
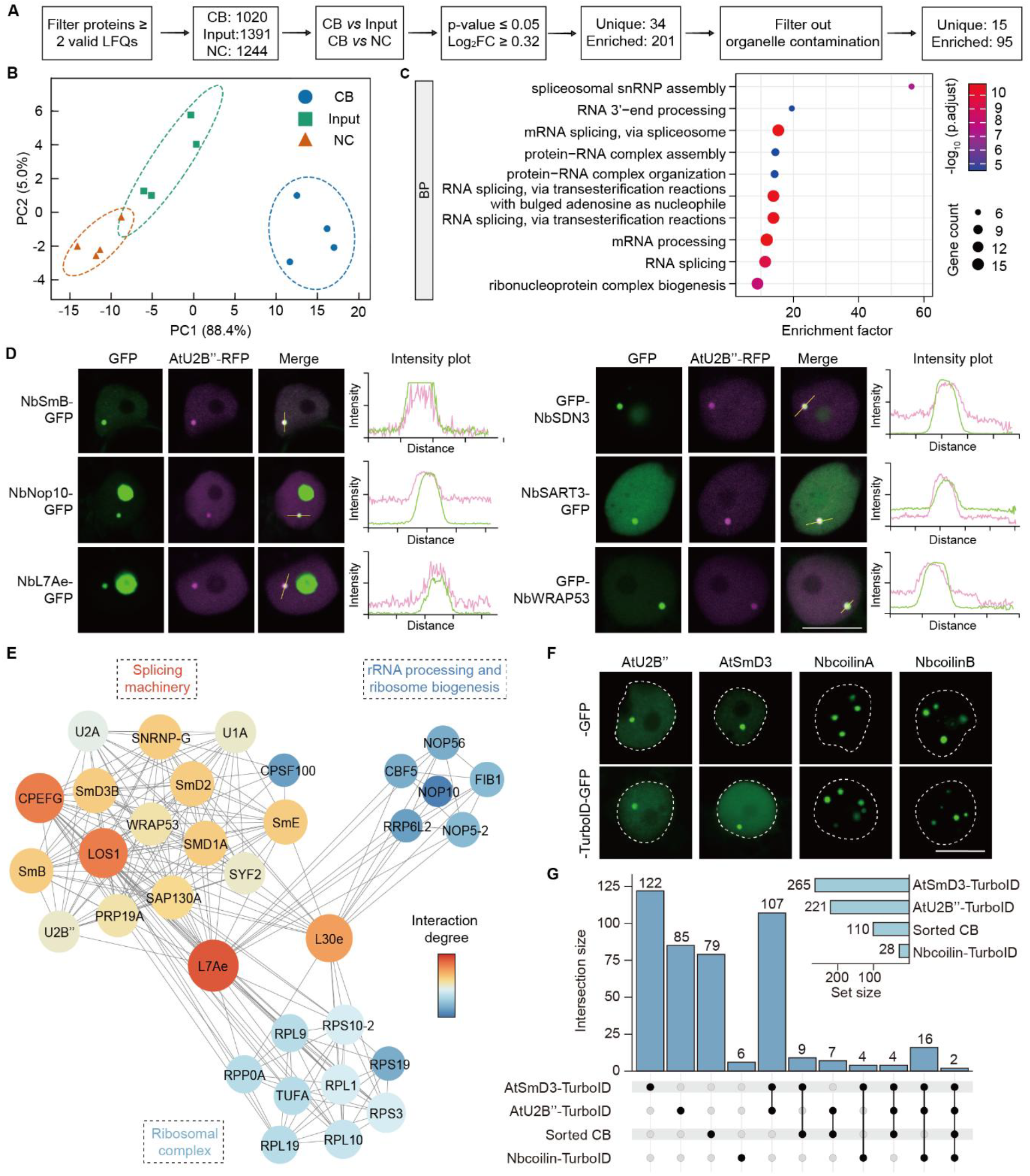
Analysis of the Cajal body proteome. A. Workflow summary of proteomic analysis to identify proteins enriched in CBs compared to input or NC. Label-free quantification (LFQ) values were used to identify proteins enriched in CB samples. B. Principal component analysis (PCA) based on LFQ intensities of proteins detected in samples of FANS-FAPS isolated Cajal bodies (CB), input, and NC. Proteins with LFQ values > 0 in at least two replicates per group were retained. Data were z-score-normalized before analysis. C. Top 10 Gene Ontology (GO) terms Biological Process (BP) enriched among FANS-FAPS isolated CB proteins. D. Representative confocal images showing localization of identified candidate proteins fused to GFP. AtU2B’’-RFP is used as CB marker. Line plots show fluorescence intensity profiles across the CB. Scale bars = 10 μm. E. Protein-protein interaction network of 110 FANS-FAPS isolated CB proteins (from A), generated using STRING and visualized in Cytoscape. Edges represent experimentally validated interactions. Disconnected nodes were hidden. F. Subcellular localization of selected CB markers fused to TurboID-GFP in *N. benthamiana* leaves at 40 hours post-agroinfiltration. Scale bar = 10 μm. G. Overlap of proteins identified by FANS-FAPS and proximity labelling assays. Data are based on three independent biological replicates.

Principal component analysis (PCA) was performed on the unfiltered proteomic data to gain an initial overview of the dataset and assess differences among sample groups. This analysis revealed clear separation between CB, input, and NC samples, each represented by four independent biological replicates (Figure 2B). The first two principal components, PC1 and PC2, accounted for 88.4% and 5.0% of the total variance, respectively. CB samples clustered distinctly from both input and NC, which showed greater similarity to each other. Pearson correlation analysis showed high intra-group correlation among the four biological replicates of each group (r = 0.94–1.00), indicating strong reproducibility (Supplementary Figure 1B). In contrast, lower inter-group correlations (r = 0.78–0.91), particularly between CB and either input or NC, further support the distinct proteomic composition of the CB compared to the nuclear fraction.

A total of 110 proteins were identified in the FANS-FAPS isolated CB-like GFP-positive particles. 15 were uniquely detected, while an additional 95 were significantly enriched compared to input or NC, with coilin being the most highly enriched (Supplementary Figure 1C and 1D; complete list available in Supplementary Table 1). “Unique proteins” might exclusively localize to the CB or be present in the total nuclear fraction in undetectable amounts. Of the 11 plant CB proteins experimentally confirmed in previous studies^8,10,12–15^, we successfully identified 9 as enriched in our dataset, namely coilin, U2B’’, U2A, U1A, SmD3, U1-70K, Nop10, fibrillarin, and AGO4, underscoring the robustness and specificity of the obtained CB proteome (Supplementary Figure 1C and 1D, Figure 3A). The two undetected plant proteins previously described as CB-localized are DCL3 and RDR2, components of the RdDM pathway like AGO4, and shown to overlap with coilin in 82 and 83% of nuclei in *Arabidopsis thaliana* protoplasts in immunolocalization experiments, suggesting a dynamic CB localization^15^. Importantly, *DCL3* and *RDR2* exhibit expression levels approximately ten times lower than *AGO4* in *N. benthamiana* leaf tissue^16^, which could explain their absence from the CB proteome obtained here. In addition, the *N. benthamiana* orthologs of several CB proteins previously characterized in mammalian systems but not yet reported in plants were also detected, including SmB, SART3, WRAP53, Nop58, Nop56, and TFII^17–21^.

**Figure 3.**
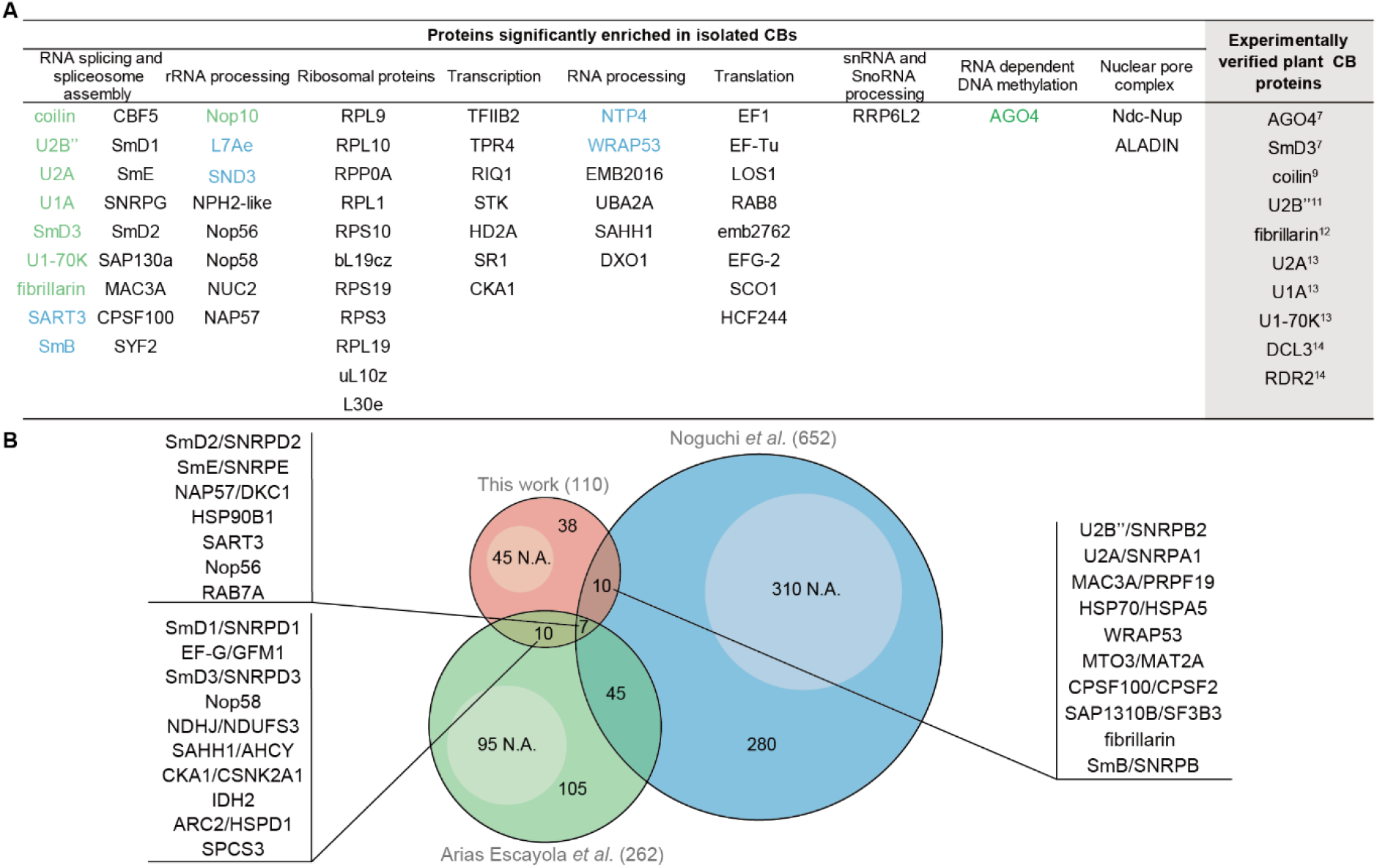
Conservation of CB proteins across kingdoms. A. Functional categorization of 60 proteins identified in FANS-FAPS isolated plant CBs. Proteins labelled in green were previously reported as plant CB proteins, while those in blue have been experimentally confirmed as CB proteins in this work. B. Comparative analysis of overlapping proteins identified across three studies: N.A. indicates no ortholog was found in *Arabidopsis thaliana* or *Homo sapiens*, respectively.

To assess the quality of our dataset, we selected eight proteins from this list for subcellular localization analysis. These included two well-characterized plant CB components, U2A and Nop10; three proteins previously reported as CB components in mammalian systems but not in plants, namely SmB, SART3, and WRAP53; and the three uncharacterized candidates SDN3, NTP4, and L7Ae. Confocal microscopy revealed that all eight proteins localized to distinct nuclear foci that overlapped with the CB marker AtU2B’’-RFP, confirming their localization to this compartment (Figure 2D and Supplementary Figure 1H).

Due to limited functional annotation of the *N. benthamiana* genome, the *A. thaliana* orthologs of identified CB proteins were used to perform functional enrichment analyses. Gene Ontology (GO) enrichment analysis using the biological process (BP) ontology revealed strong enrichment for terms such as “spliceosome assembly”, “RNA splicing,” and “ribonucleoprotein complex assembly” (Figure 2C), in line with the known functions of the CB. Overrepresented cellular component (CC) terms were topped by “Cajal body” and “snRNP complex”, while those in the molecular function (MF) ontology included “snRNA binding” and “snoRNA binding” as the most highly enriched (Supplementary Figures 1D and 1E). In contrast, proteins predominantly enriched in the input and NC fractions were associated with more general nuclear functions (full list available in Supplementary Table 2), with a significant enrichment for “nucleocytoplasmic transport,” “gene expression,” and “chromatin organization” (BP), “nucleoplasm,” “chromatin,” and “chromosome” (CC), and “RNA helicase activity” and “chromatin binding” (MF) (Supplementary Figure 1F). These findings highlight a clear functional distinction between the nuclear background and the CB-specific proteome. In agreement with the results of the functional enrichment analyses, protein interaction network analysis on identified CB proteins, based on experimentally validated interactions from the STRING database^22^ revealed three main protein clusters, related to the splicing machinery, ribosomal complexes, and rRNA processing, respectively (Figure 2E), with two ribosomal proteins, L7Ae and L30e, connecting the three groups.

Given the previous use of PL techniques to profile proximal proteomes within biomolecular condensates^4–6^, we performed TurboID-based PLs to complement the proteomic dataset obtained from FANS-FAPS isolated CBs. NbcoilinA, a CB scaffold protein used in other CB-proximity labelling studies^4–6^, and its paralog NbcoilinB were used as baits. Two additional characterized plant CB proteins involved in distinct small nuclear ribonucleoprotein complexes, AtU2B″, a component of the U2 snRNP^23^, and AtSmD3, a member of the Sm core^24^, were also included. Each protein was fused to either GFP or GFP-TurboID to assess the localization of the resulting proteins. Both GFP and GFP-TurboID fusion proteins localized to CBs (Figure 2F). Notably, when expressed under the strong 35S promoter, both NbcoilinA-GFP/GFP-TurboID and NbcoilinB-GFP/GFP-TurboID formed multiple nuclear foci yet maintained highly specific localization with no detectable signal in the nucleoplasm, suggesting that these scaffolding proteins are limiting factors for CB assembly. Proximity labelling assays were performed using TurboID-tagged, GFP-free versions of the four bait proteins. Compared to AtU2B’’ and AtSmD3, NbcoilinA and NbcoilinB exhibited lower accumulation levels; nevertheless, all fusion proteins displayed biotinylation activity (Supplementary figure 2A). PL identified 265 proteins labelled by AtSmD3-TurboID, 221 by AtU2B’’-TurboID, and 28 by Nbcoilin-TurboID (NbcoilinA + NbcoilinB) (Figure 2G, full list available in Supplementary Table 3). Functional enrichment analysis revealed distinct profiles (Supplementary figure 2B-D): proteins labelled by AtU2B″-TurboID were enriched in “ribosomal RNA processing” and “RNA splicing” (BP), “ribosomal structural constituents” (MF), and “ribosomal protein” (CC); AtSmD3-TurboID-labelled proteins showed enrichment in “mRNA processing” and “non-membrane organelle assembly” (BP), “single-stranded RNA binding” (MF), and “ribosomal protein” (CC); Nbcoilin-TurboID-labelled proteins were associated with “centromere complex assembly” (BP), “structural molecule activity” (MF), and “ribosomal protein” (CC). These results highlight the bait-dependent nature of proximal proteomes, despite all baits localizing to the same MLO. A total of 22 proteins overlapped between the proximity-labelling datasets and the FNAS-FAPS isolated CB proteome (Figure 2F). GO enrichment analysis of these proteins revealed a significant overrepresentation of “splicing” (BP), “poly(A) RNA binding” (MF), and “nuclear body” (CC) categories (Supplementary figure 2E).

A significant overlap was observed between the AtSmD3- and AtU2B’’-TurboID datasets, with 107 shared proteins. GO analysis of these overlapping proteins (Supplementary table 6) indicated an enrichment in general nuclear functions such as “translation” and “gene expression” (BP). Although 13 of these 107 proteins were associated with splicing, a known function of CB^7,25^, the overall profile suggested a broader nuclear labelling pattern. This broader labelling may be attributed to the observed partial localization of AtU2B’’ and AtSmD3 in the nucleoplasm (Figure 2E), which could allow labelling of non-CB-proximal proteins.

We next compared the FANS-FAPS isolated plant CB proteome to the human cell CB proximity proteomes produced by Arias Escayola *et al*. (2025) and Noguchi *et al*. (2024) (Figure 3B). Orthologous proteins were identified through reciprocal BLAST searches against the *A. thaliana* or *Homo sapiens* proteomes using criteria of pairwise identity ≥30%, E-value ≤10−^10^, and query coverage ≥50%. Conserved CB proteins across kingdoms include Sm proteins, U2 RNPs, Nop56/58, fibrillarin, SART3, NAP57, and WRAP53. However, many CB proteins were found to be kingdom-specific. Among the 110 proteins identified in this study, 45 lacked orthologs in humans, including the experimentally validated NPT4 and SDN3, which play a role in RNA processing. Similarly, 310 out of 652 proteins identified by Noguchi *et al*. (2024) and 95 out of 262 proteins identified by Arias Escayola *et al*. (2025) lack orthologs in *A. thaliana*. Among them, TOE1^4^, IR2BP1^4^, TDP-43^5^ and RIF1^5^ were experimentally confirmed as CB-localized. While telomere synthesis-associated proteins, such as dyskerin (DKC1) and its plant ortholog NAP57, are conserved across kingdoms, other core animal telomere-associated proteins like TERT^26^ do not have orthologs in plants. These observations underscore that despite functional conservation of the CB, the specific molecular machineries involved can differ substantially across kingdoms, reflecting specialized adaptations to unique cellular and physiological contexts of each lineage.

The FANS-FAPS isolation method developed in this study shows strong potential for broader applicability across diverse organisms and MLOs, both in plants and animals. The foundational principle, using fluorescently tagged markers coupled with FAPS, is adaptable by simply switching the marker protein. Optimization of buffer compositions and sorting conditions is likely to allow the effective application of this method to a diversity of cells and condensates.

To conduct an effective MLO isolation through FAPS, the selection of the sorting marker protein is a crucial step. Coilin was the marker of choice for CB isolation because it structurally defines this compartment and displays a highly specific localization. Previous studies demonstrated that coilin knock-out or knock-down directly results in the reduction of CBs in both animals and plants^10,27–29^ and significantly reduces cellular accumulation of other CB proteins, such as AGO4^30^ or causes mislocalization of other components like U-RNPs^31^, emphasizing coilin’s essential role in CB formation and maintenance. An ideal MLO marker should be specifically localized to the target compartment and functionally involved in its assembly or activity, ensuring that the isolated particles represent *bona fide* and biologically relevant organelles.

PL has emerged as a powerful tool for analyzing partial MLO proteomes by using engineered enzymes, such as BioID, APEX, or TurboID, fused to known condensate-resident proteins. Limitations of this approach include a small effective labelling radius (10 nm) compared to typical MLO dimensions (0.1 –5 µm); additionally, the water-soluble nature of biotin may hinder effective diffusion into dense, phase-separated MLOs. In this work, PL of CB using different resident proteins as baits revealed limited overlap among datasets, highlighting the strong dependence of proteomic outputs on the choice of marker and the partial nature of the resulting proteomes. In contrast, the CB proteome obtained via FANS-FAPS represents the most comprehensive dataset, showing no signs of nucleoplasmic contamination, as evidenced by strong overlap with known plant CB proteins, localization studies, and functional GO analysis. These results suggest that direct isolation of intact MLOs enables a more complete and unbiased proteomic profiling, offering a superior alternative to marker-dependent PL approaches. Nevertheless, combining PL with direct isolation of MLOs may further enhance the specificity and resolution of proteome characterization.

Understanding the dynamics of CBs and other nuclear MLOs is essential for elucidating their biological functions. CBs serve as organized platforms for enzymes and substrates, significantly enhancing specific cellular processes, such as RNA processing, ribosome biogenesis, and telomere maintenance^7,9,25^. Mathematical modelling suggests that the presence of four CBs per nucleus could increase snRNP folding efficiency 11-fold^32^. The number and size of CBs vary significantly among cell types, developmental stages, and stress conditions. For instance, variations occur between vegetative and sperm cells in *A. thaliana* pollen^33^. Viral infections such as groundnut rosette virus (GRV) infection can induce CB restructuring into multiple CB-like bodies^34^, demonstrating CB plasticity and responsiveness to environmental stimuli, similar to other MLOs such as the nucleolus^35^. The dynamic, membraneless nature of MLOs enable rapid exchange and re-distribution of components in response to environmental or developmental cues. Shared localization of proteins such as AGO4 (CB and AB-body)^8^ and fibrillarin (CB and nucleolus)^34^ exemplifies this plasticity. Notably, under viral stress, only CB-localized AGO4 interacts with V2 from tomato yellow leaf curl virus (TYLCV), which suppresses the RdDM-mediated viral genome methylation^36^. Similarly, GRV ORF3 reorganizes CBs, causing fusion with nucleoli for viral movement. These findings point to the potential significance of quantitative proteomic changes in CBs across physiological states. By leveraging the ability to isolate intact CBs and systematically analyze their proteomes, it is now possible to investigate how their composition dynamically responds to developmental cues and environmental stress. Concurrent isolation and analysis of other nuclear MLOs will enable comparative approaches, providing insights into the functional interplay between CBs and other nuclear compartments. This, in turn, will advance our understanding of how CBs contribute to nuclear organization and function under diverse physiological contexts. Moreover, such studies will shed light on the biophysical and molecular principles governing MLO formation and the dynamic exchange of their components— mechanisms that underpin their plasticity and biological relevance.

## METHODS

### Plant material and growth conditions

*Nicotiana benthamiana* plants were grown in a controlled growth room under long-day conditions (16 h light and 8 h dark) at 25°C.

### Plasmids construction

All primers used in this study are summarized in Table S7.

For construction of the CB marker used for FAPS, the *pRPS5A* promoter was adapted from the GreenGate system^11^, with modifications to the overhang sequences. The *pRPS5A* promoter and *NbcoilinA* (Niben101Scf04802g01014) were assembled into Level II vector from the GoldenGate-based toolkit^37^ via *Bsa*I cut-ligation, along with the following modules: DE-GFP, EF-nosT, and FG-dummy. *N. benthamiana* genes *NbU2A* (Niben101Scf02581g05011), *NbSmB* (Niben101Scf01968), *NbNop10* (Niben101Scf02243g01029), *NbL7A*e(Niben101Scf03147g07006), *NbSDN3* (Niben101Scf12361g03004), *NbSART3* (Niben101Scf00705g05008), *NbNTP4* (Niben101Scf03208g00007), and *NbWRAP53* (Niben101Scf07372g00012), were cloned into Level II vector at the CD position using *Bsa*I/Esp31 cut-ligation, together with modules AC-p35S, DE-GFP, EF-nosT, and FG-dummy. Gateway-compatible constructs for *AtU2B*’’ (At2g30260), *AtSmD3* (At1g76300), *NbcoilinA*, and *NbcoilinB* (Niben101Scf04939g02015) were generated by cloning into the destination vectors pGW-GOI-TurboID and pGW-GOI-TurboID-GFP^38^.

### Agrobacterium-mediated transient expression

*Agrobacterium tumefaciens* strain GV3101 was used for delivery of plasmid constructs into *N. benthamiana* leaves as previously described^38^.

### Cajal body (CB) isolation and fluorescence-activated particle sorting

Leaves from four-week-old *N. benthamiana* plants transiently transformed with pRBS5a:*NbcoilinA-GFP* (NbcoilinA-GFP) or an empty vector (NC) were crosslinked in a fixation buffer containing 10 mM Tris-HCl, 10 mM EDTA, 100 mM NaCl, and 0.1% Triton X-100, supplemented with 1% formaldehyde. Vacuum infiltration (50 psi) was applied for 10 minutes twice. Crosslinking was quenched by vacuum infiltration with 125 mM glycine for 10 minutes. Tissue was then homogenized using a blender (SEVERIN, UZ3861) in lysis buffer (15 mM Tris-HCl, 2 mM EDTA, 80 mM KCl, 20 mM NaCl, 0.5 mM spermine, 15 mM dithiothreitol, 10 mM protease inhibitor, and 0.1% Triton X-100) at 260 W in 30-second on/off mode for a total of five cycles. The homogenate was centrifuged at 800 × *g* for 10 minutes at 4 °C. The pellet was then resuspended in suspension buffer (20 mM Tris-HCl, 1.5 mM MgCl_2_, 10 mM KCl, 15 mM dithiothreitol, and 10 mM protease inhibitor for nuclei sorting (see below). Sorted nuclei were sonicated using a SONOPULS Ultrasonic Homogenizer (BANDELIN) at 40% power in 15-second on/off cycles six times. The lysate was centrifuged at 800 × *g* for 10 minutes at 4 °C, and the resulting supernatant containing released CBs was subjected to CB sorting (see below). Nuclear fractions were collected at this step from both NbcoilinA-GFP-expressing samples (Input) and EV-expressing samples (NC), then freeze-dried using an Alpha 1–4 LSC vacuum dryer (CHRIST) at –60 °C for 48 hours. The resulting powder was stored at room temperature until further use. Sorted CBs were pelleted by ultra-centrifugation at 42,000 × g for 45 minutes. The pellet was stored at –80 °C until further use.

Nuclei and CB sorting were performed using a MoFlo XDP cell sorter (Beckman Coulter) equipped with a 70 µm nozzle, operating at a sample/sheath pressure differential of 61/60 psi. Phosphate-buffered saline (PBS, pH 7.0) was used as the sheath fluid, and sorting was conducted in Purify 1 Drop mode to ensure high purity. For nuclei sorting, 0.1 µg/mL DAPI (4’,6-diamidino-2-phenylindole; Invitrogen) was used to stain nuclei and excited with a 15 mW, 375 nm laser in spherical focus mode. DAPI emission was collected using a 405/30 nm bandpass filter. GFP and side scatter were excited with a 75 mW, 488 nm laser using a cross-cylindrical focus. GFP emission was collected with a 534/30 nm bandpass filter, and side scatter with a 488/10 nm filter. Nuclei were identified as bright DAPI-positive populations compared to side scatter. Doublets and clumps were excluded by comparing DAPI peak height to area values. GFP-positive nuclei were confirmed by plotting DAPI versus GFP fluorescence and comparing to untransformed negative controls. CBs were identified as small, GFP-positive particles distinguishable from nuclei by their lower side scatter profile. Gating was optimized by comparing GFP versus side scatter plots between samples and negative controls. To ensure precise gating during sorting, live acquisition was monitored by displaying the most recent 25,000 events.

### TurboID-based proximity labelling assays

Leaves from four-week-old *N. benthamiana* plants transiently transformed with constructs to express TurboID-fused proteins or free TurboID were processed for proximity labelling assays as described^38^ and subjected to LC-MS/MS analyses.

### Western blotting

Western blot analysis was performed as previously described^38^. The primary and secondary antibodies used are the following: goat anti-GFP (Sicgen, Catalog number AB0020-500, 1:15,000 dilution); mouse anti-HA (Sigma-Aldrich, H3663; 1:3,000); mouse anti-Actin (Agrisera, AS21-461510, 1:2,000); rabbit anti-BiP (Agrisera, AS09-481, 1:2,000); Rabbit anti-H3 (Agrisera, AS09-481, 1:5,000); anti-mouse peroxidase (Sigma-Aldrich, A2554; 1:15,000 dilution); anti-goat peroxidase (Sigma-Aldrich, A8919, 1:20,000); anti-rabbit peroxidase (Sigma-Aldrich, A0545, 1:15,000).

### Liquid chromatography – tandem mass spectrometry (LC-MS/MS) analysis

Pellets collected after CB isolation were resuspended in 4x NuPAGE LDS sample buffer and heated at 95 °C for 10 minutes to reverse formaldehyde crosslinking prior to SDS-PAGE. In four independent biological replicates, 7.65 million, 8.9 million, 8.4 million, and 9.1 million particles were collected, respectively. Proteins were separated by SDS-PAGE, followed by in-gel digestion^39^. Extracted peptides were desalted using C18 StageTips^40^. Equal peptide amounts were subjected to LC-MS analysis on a VanquishNeo nano-UHPLC coupled to Exploris 480 mass spectrometer via nanoelectrospray ion source (Thermo Fisher Scientific). Peptides were separated on a custom-packed nano-LC column (20 cm length, 75 µm inner diameter) containing 1.9 µm C18 ReproSil-Pur beads using a 49-minute segmented gradient from 4% to 55% of solvent B (80% acetonitrile in 0.1% formic acid) at a flow rate of 300 nL/min. MS and MS/MS spectra were acquired in positive ion mode with a resolution of 60,000 for MS1 and 30,000 for MS2 scans. The AGC target was set to “standard,” and the maximum injection time was set to “auto.” Precursor ion selection was performed using a 1.3 Th isolation window, and the top 20 most intense precursors were selected for higher-energy collisional dissociation (HCD) per scan cycle. Dynamic exclusion was set to 30 seconds. TurboID samples were analyzed on an Easy-nanoLC coupled to a Q Exactive HF mass spectrometer (Thermo Fisher Scientific) using a 49-min gradient (10–33–50–90% solvent B) at 200 nL/min. The top seven precursors were fragmented, with resolution set to 60,000 for both MS and MS/MS spectra and an AGC target of 3E6.

### MS data processing and statistical analysis

Raw mass spectrometry (MS) data were analyzed using MaxQuant software (version 2.2.0.0)^41^, employing the integrated Andromeda search engine^42^. Spectra were searched against a *N. benthamiana* protein Niben101 database (Sol Genomics Network), supplemented with a database of 286 common laboratory contaminants. Sequences for NbcoilinA-GFP were added as custom entries to the search database. Default MaxQuant parameters were used, with intensity-based absolute quantification (iBAQ) and label-free quantification (LFQ) enabled. The “match between runs” feature was activated for samples within the same experimental group to improve peptide identification consistency.

Statistical comparisons were conducted in Perseus software (version 1.6.15.0) following log_10_ transformation of LFQ intensity values. Significance testing was performed using a two-sample t-test with permutation-based false discovery rate (FDR) control set to 0.05 and S_0_ = 0. Data were filtered to retain only proteins with at least two valid LFQ values in one of the comparison groups. The Niben101 ID was converted to TAIR10 ID at Niben2Arab.

## Confocal laser scanning microscopy

Confocal imaging of the subcellular localization of GFP-or RFP-tagged proteins expressed in *N. benthamiana* epidermal cells was performed using a Zeiss LSM880 confocal laser scanning microscope. GFP was excited with a 488 nm laser and detected between 500–550 nm. RFP was excited with a 561 nm laser and detected between 570–630 nm. For nuclear staining, 0.1 µg/mL DAPI was applied and excited using a 405 nm laser, detected in the 420–470 nm range.

## DATA AVAILABILITY

The mass spectrometry proteomics data have been deposited to the ProteomeXchange Consortium via the PRIDE^43^ partner repository with the dataset identifier PXD065938.

## ACKNOWLEDGEMENTS

The authors thank the GeminiTeam Lab for the critical reading of the manuscript, with especial thanks to Bettina Stadelhofer and Ying Li for their excellence technical support. We also thank Richard Kormelink and Mandy Ravensbergen for fruitful discussions. We are indebted to the Central Facilities at ZMBP, particularly FACS, Microscopy, and Plant Cultivation, for their outstanding technical assistance. This work is funded by the Excellence Strategy of the German Federal and State Governments, the ERC Consolidator Grant GemOmics (101044142), and a Promotion of Junior Researchers grant to YZ (PRO-ZHOU-2023-20) from the University of Tübingen.

## AUTHOR CONTRIBUTIONS

RL-D, YZ and KB conceived the project; YZ, HT, AV-L, MW, LW, MS, ID-B, BM and KB performed the experiments and analyzed data; YZ and HT prepared the figures; YZ and RL-D wrote the manuscript, with input from all authors.

## SUPPLEMENTARY MATERIAL

## SUPPLEMENTARY FIGURES

**Supplementary figure 1. Validation and functional enrichment analysis of the sorted Cajal body (CB) proteome**.

A. Gating strategy for isolating GFP-positive nuclei using Fluorescence activated nuclei sorting (FANS). H: height; A: area.

B. Pearson correlation matrix of LFQ intensities among biological replicates of CB, input, and NC samples, showing high reproducibility within each sample group. r > 0.9: reproducible technical or biological replicates; r ∼ 0.6–0.8: related but distinct interaction profiles.

C, D: Volcano plots illustrating the differential enrichment of proteins in the FANS-FAPS isolated CB compared to either the input (C) or the NC (D). Proteins with a two-tailed Student’s t-test p < 0.05 and a fold change greater than 1.25 were considered significantly enriched in the CB.

E. Top 10 Gene Ontology (GO) terms Cellular component (CC) enriched among isolated CB proteins.

F. Top 10 Gene Ontology (GO) terms Molecule function (MF) enriched among isolated CB proteins.

G. GO enrichment analysis of enriched proteins shared between input and NC samples, across biological process (BP), molecular function (MF), and cellular component (CC) categories.

H. Representative confocal images showing localization of identified candidate proteins fused to GFP to the CB. AtU2B’’-RFP is used as CB marker. Line plots show fluorescence intensity profiles across the CB Scale bars = 10 μm.

**Supplementary figure 2. TurboID-based proximity labelling assays with CB markers and functional enrichment analysis of identified proteins**.

A. TurboID-fused CB markers retain biotinylation activity in *N. benthamiana* leaves in the presence of 50 μM exogenous biotin. Western blot analysis shows accumulation of the fusion proteins (α-HA) and streptavidin-HRP detection of biotinylated proteins from samples expressing AtU2B’’-TurboID (61 kDa), AtSmD3-TurboID (49 kDa), NbcoilinA-TurboID (126 kDa), NbcoilinB-TurboID (124 kDa) and TurboID alone (35 kDa). The corresponding fusion protein bands are indicated by red asterisks.

B–E. Gene Ontology (GO) enrichment analyses of proteins identified in TurboID-based proximity labelling assays with the CB markers in A. B, AtU2B’’-TurboID; C, AtSmD3-TurboID; D, Nbcoilin-TurboID; E, 22 proteins overlapping between FANS-FAPS isolated CB and TurboID-based proximity labelling. Enrichment results are shown for Biological Process (BP), Molecular Function (MF), and Cellular Component (CC) categories.

## SUPPLEMENTARY TABLES

**Supplementary table 1**. List of putative plant Cajal body components identified by FANS-FAPS CB isolation.

**Supplementary table 2**. Proteins enriched in input and NC nuclear fractions.

**Supplementary table 3**. Proteins labelled by AtU2B’’-TurboID, AtSmD3-TurboID, and Nbcoilin-TurboID, compared to TurboID-only controls.

**Supplementary table 4**. Gene ontology analysis of putative Cajal body components (from Supplementary Table 1) and proteins enriched in the nuclear fraction (from Supplementary table 2).

**Supplementary table 5**. Gene ontology analysis of proteins labelled by AtU2B’’-TurboID, AtSmD3-TurboID, and Nbcoilin-TurboID (from Supplementary Table 3).

**Supplementary table 6**: Gene ontology analysis of proteins overlapping between the indicated datasets.

**Supplementary table 7**. Primers used in this work.

**Supplementary table 8**. Cross-kingdom identification of orthologous proteins.

