## Supplementary figures for "Proteome profiling of plant Cajal bodies uncovers kingdom-specific components"

Supplementary figures 1-2

### SUPPLEMENTARY TABLES

Supplementary tables 1-8

### SUPPLEMENTARY FIGURES

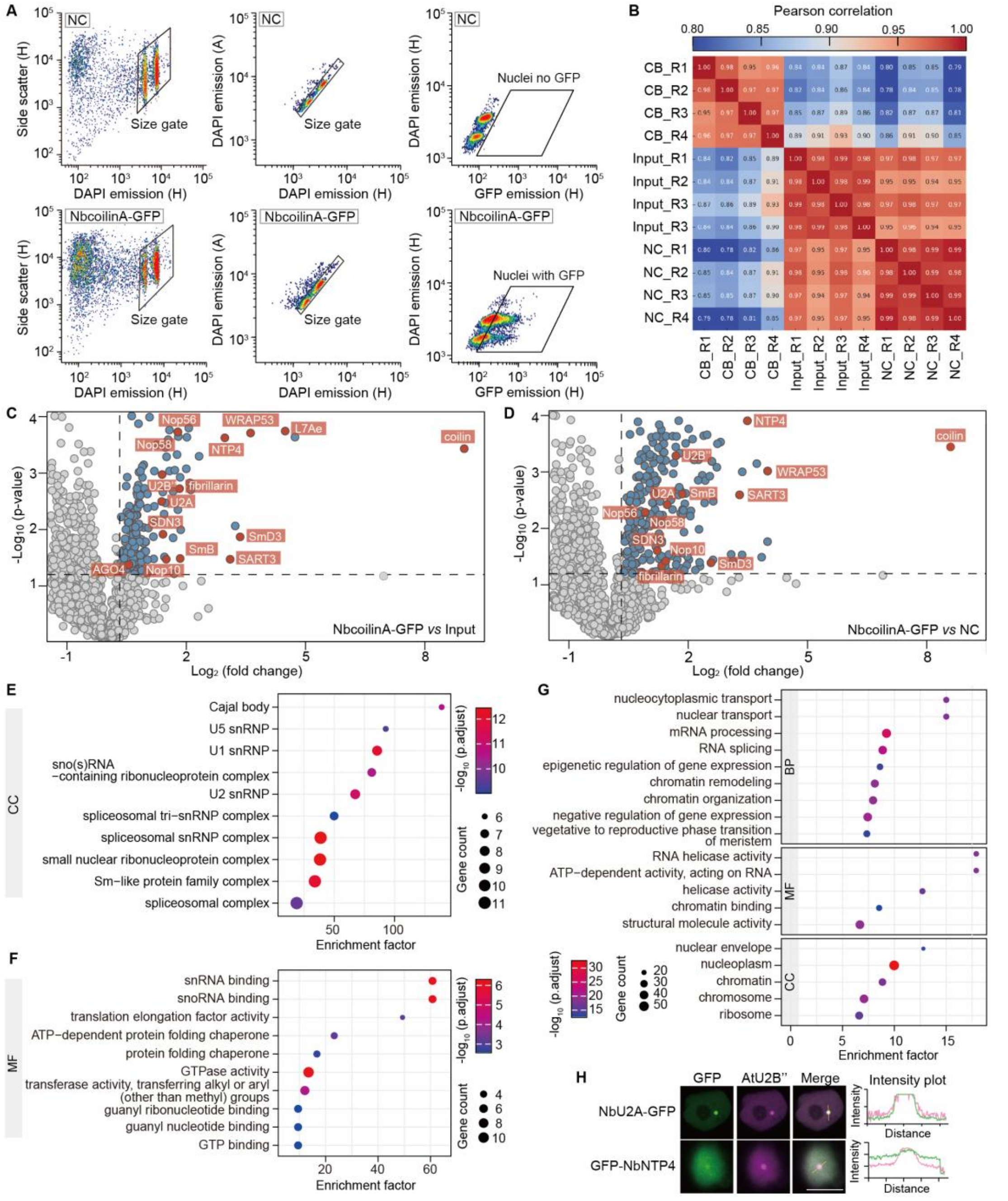


**Supplementary figure 1. Validation and functional enrichment analysis of the sorted Cajal body (CB) proteome.**

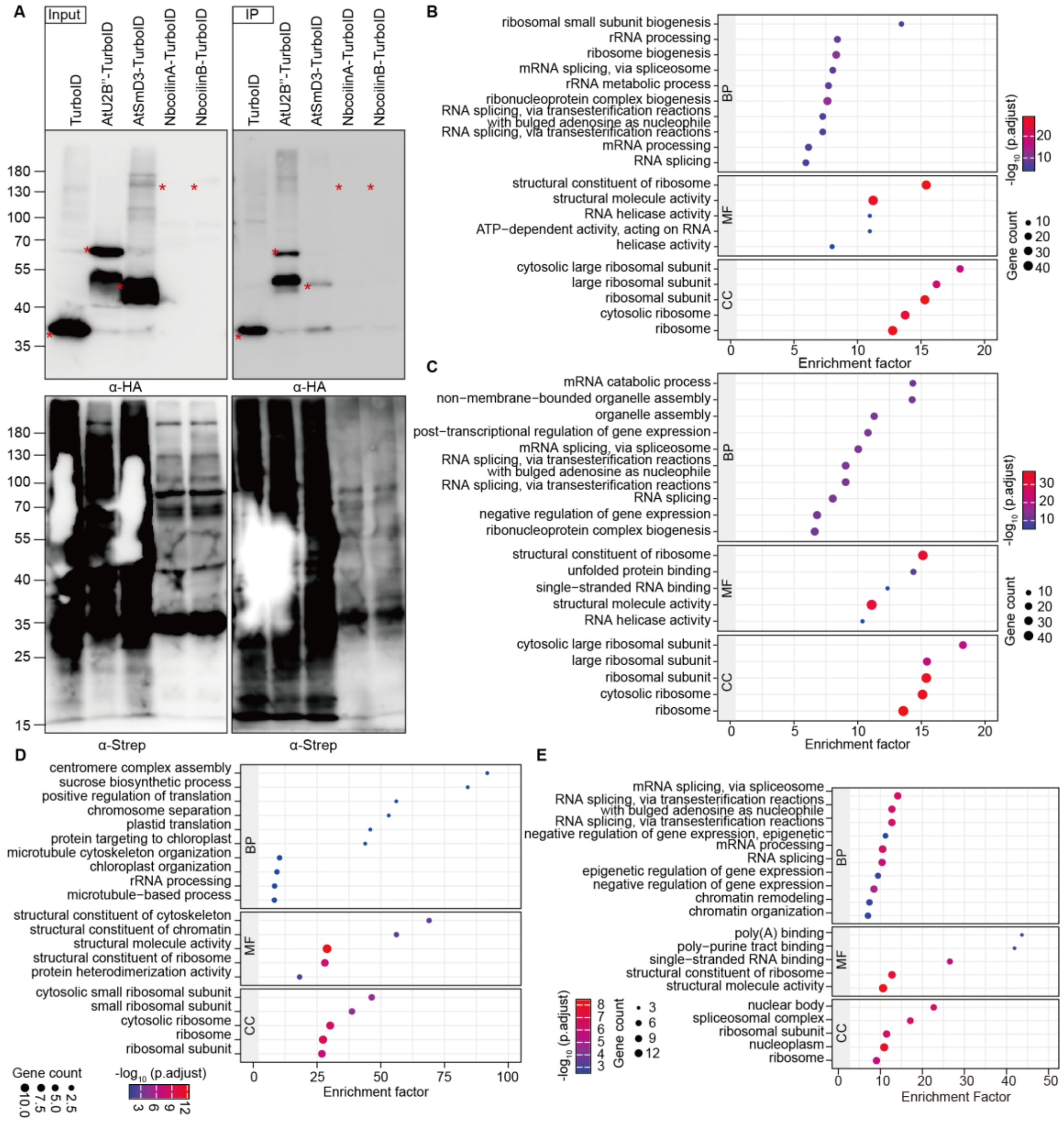


**Supplementary figure 2. TurboID-based proximity labelling assays with CB markers and functional enrichment analysis of identified proteins.**
